# Mapping the neural circuitry of predator fear in the nonhuman primate

**DOI:** 10.1101/2020.04.24.057869

**Authors:** Quentin Montardy, William C. Kwan, Inaki C. Mundinano, Dylan M. Fox, Liping Wang, Cornelius T. Gross, James A. Bourne

**Affiliations:** Shenzhen Key Lab of Neuropsychiatric Modulation and Collaborative Innovation Center for Brain Science, Guangdong Provincial Key Laboratory of Brain Connectome and Behavior, CAS Center for Excellence in Brain Science and Intelligence Technology, Brain Cognition and Brain Disease Institute (BCBDI), Shenzhen Institutes of Advanced Technology, Chinese Academy of Sciences, Shenzhen-Hong Kong Institute of Brain Science-Shenzhen Fundamental Research Institutions, Shenzhen 518055, China; Australian Regenerative Medicine Institute, Monash University, Clayton, VIC 3800, Australia; Epigenetics & Neurobiology Unit, EMBL Rome, European Molecular Biology Laboratory, Via Ramarini 32, 00015 Monterotondo (RM), Italy

## Abstract

In rodents, innate and learned fear of predators depends on the medial hypothalamic defensive system, a conserved brain network that lies downstream of the amygdala. However, it remains unknown whether this network is involved in primate fear. Here we demonstrate that visually evoked predator fear recruits a homologous medial hypothalamic defense system in the nonhuman primate

## Introduction

Lesions of the amygdala block the processing of learned and innate fear stimuli in multiple species, including humans (1–4). However, fear induced by internally generated stimuli, such as the inhalation of carbon dioxide, do not require the amygdala (5). This observation suggests that circuits downstream of the amygdala are sufficient to sustain the behavioral and emotional correlates of fear. This view is supported by extensive work in rodents demonstrating that a circuit from the amygdala to the medial hypothalamus and brainstem, called the medial hypothalamic defensive system, is both necessary and sufficient for innate and learned defensive responses to predators (6–10). Furthermore, in humans, direct electrical stimulation of the ventromedial hypothalamus (VMH) is sufficient to elicit feelings of intense fear and trigger panic attacks, suggesting that the medial hypothalamus may also participate in human fear (11). Unlike rodents, however, primates typically depend exclusively on visual cues to detect and respond to innate threat cues, yet it remains unknown whether such stimuli are sufficient to recruit the medial hypothalamic defensive system and by what pathway this might occur.

Here we developed a method to elicit robust predator-evoked escape responses in a nonhuman primate (marmoset monkey) under controlled laboratory conditions, and used c-Fos mapping combined with anatomical tract tracing to determine the neural circuits involved. The common marmoset, *Callithrix jacchus*, is an appealing primate species to study the link between neural circuits and behavior because of its small size and its rich repertoire of affiliative and agonistic social behaviors (12). The defensive responses of marmosets to predators in the wild have been described and include visual scanning, alarm calling, mobbing, avoidance, freezing, and flight (13,14). Exposure to toy snakes, cats or raptors have been used in the laboratory setting to induce visual scanning, alarm calling, freezing and threat displays (15–17). However, to the best of our knowledge, the full repertoire of active defensive responses, including flight and post-flight vigilance observed in response to predators in the wild, has not been reported under laboratory conditions.

## Results

### Snake presentation evokes flight, sustained avoidance, and vigilance in the primate

Animals were pre-trained over a two-month period to enter a transparent plexiglass transport box juxtaposing the home cage. This was followed by 2-4 weeks of habituation to a testing room, during which time the animal was given access via a small door in the transport box to an opaque nest box (**Fig. 1AB**). Following habituation, the animals underwent surgery for injection of fluorescent anterograde (dextran amine) and retrograde (cholera toxin B) tracers into the VMH (**Fig. 2B**). Recent advances in MRI-guided stereotaxic surgery in the marmoset(18) have enabled the precise delivery of reagents to discrete brain regions. Following recovery, the animals were trained for an additional five days, during which time their baseline behaviors in the isolated testing room were recorded. On the following day, a remotely activated canopy positioned in front of the transport box was raised to reveal the threat stimulus – a moving rubber snake, while control animals were exposed to a neutral stimulus – a small black box (**Fig. 1B**), for a continuous duration of 45 minutes (**Fig S1A**). Exposure to the threat stimulus elicited robust defensive responses in all animals, consisting of flight to the rear of the cage and hypervigilance, and culminating in escape to the adjacent nest box (**Fig. 1BC**, **Video S1**). The behavioral state of the animal was scored using a Defensive Behavior Index, consisting of a weighted sum of distance from the stimulus (from −0.8 when near the stimulus to +1 when in the retreat box) and defensive behavior (from −1 when grooming to +2 during flight; **Fig. 1C**). As long as the threat stimulus was visible, animals remained principally in the nest box (**Fig. 1D**, Pre = 4.3% vs Threat = 93%, P < 0.0001, t = 14.69, df = 8), showing a significant reduction in exploration (**Fig. 1E**, Pre = 6.0 vs Threat = 1.2, P < 0.05, t = 2.75, df = 8) and increase in lurking (**Fig. 1F**, Pre = 0.0 vs Threat = 2.1, P < 0.01, t = 4.75, df = 8) and staring (**Fig. 1G**, Pre = 0.04 vs Threat = 1.2, P < 0.05, t = 2.78, df = 8). Scanning, on the other hand, was not significantly affected by threat presentation (**Fig. 1H**, Pre = 0.48 vs Threat = 0.39). In comparison, animals exposed to the control stimulus demonstrated no signs of startle or flight (**Fig. S1B**, **Video S2**), and mainly remained outside of the retreat box with no change in exploration (**Fig. S1B-E**). On the contrary, control animals stood closer to the front of the transport box to observe the stimulus (**Fig. S1F**, Pre = 0.1 vs Stim = 3.0, P < 0.01, t = 29, df = 2), while scanning behaviors remained unchanged (**Fig. S1G**). In a subset of animals (N = 3) the threat stimulus was presented for five minutes and then hidden again under the canopy to assess recovery (**Fig. S2A**). Following removal of the threat stimulus, animals exited the nest box and their Defensive Behavior Index returned to baseline (latency = 43-120 sec, **Fig. S2B,C**). During this recovery period, animals spent significantly less time in the retreat box (**Fig. S2D**, Threat = 90% vs Recovery = 38%, P < 0.05, t = 3.67, df = 4), but other behaviors such as lurking, staring and scanning did not return to baseline levels (**Fig. S2E-H**). A subset of animals was exposed for 45 minutes to either the threat (N = 2) or neutral stimulus (N = 2) and processed for histological analysis and cFos expression (**Fig. 1A, Fig. S1A**).

**Figure 1.**
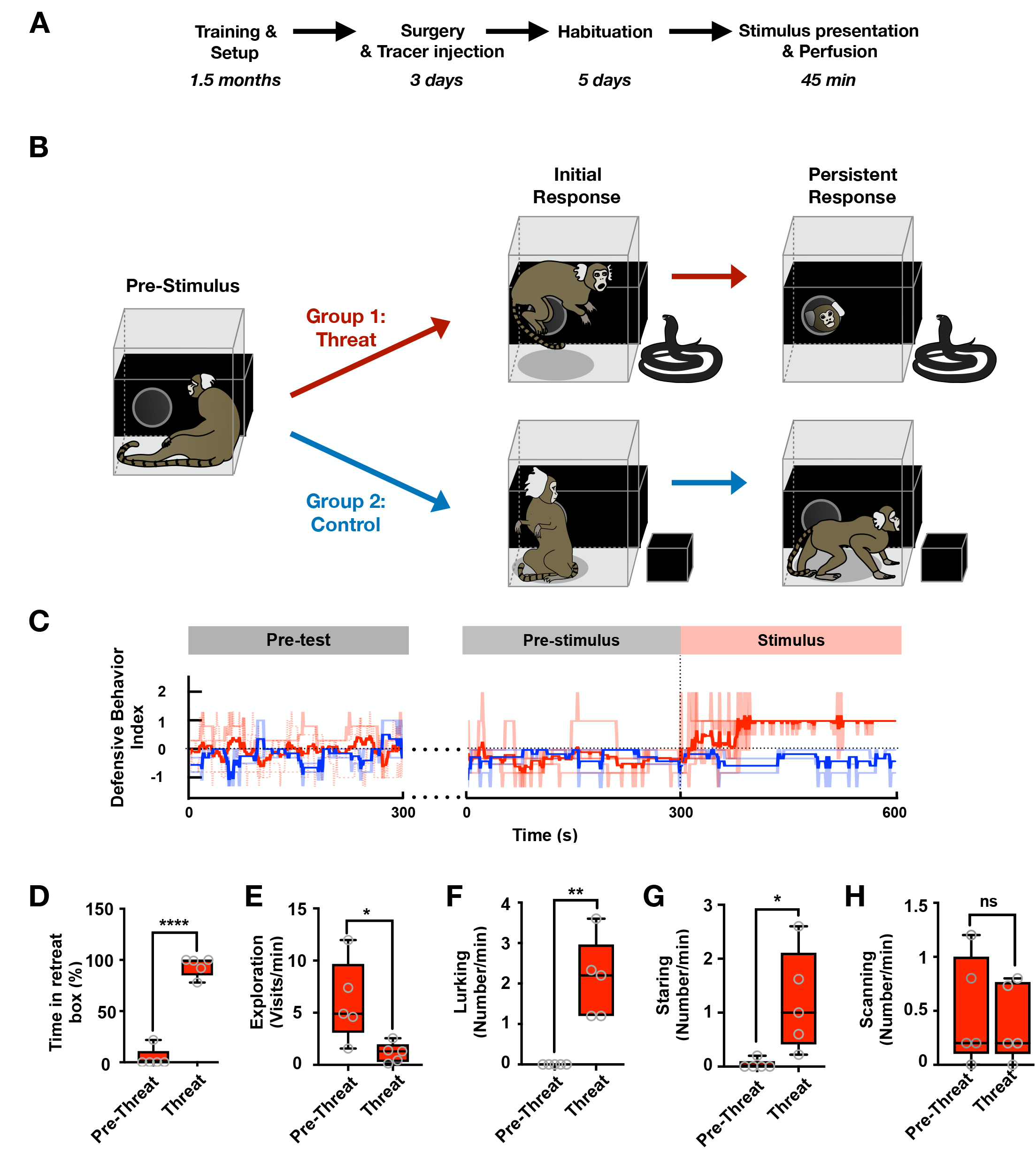
Snake presentation evokes flight, sustained avoidance, and vigilance in the primate. **(A)** Following an extensive period of habituation training animals underwent surgery for the local delivery of anterograde and retrograde tracers in VMH. After recovery training was continued for 5 days before exposing the animal to the experimental stimulus. Forty-five minutes after stimulus exposure the animal was anesthetized, perfused, and its brain prepared for cFos immunostaining. Animals were randomly assigned to groups exposed to either an animated rubber snake (threat) or a black cardboard box (control). Images indicate representative behaviors evoked by the stimuli during the initial and persistent response phases of the test. (**C**) Defensive Behavior Index (light color, individual traces; dark color, mean; 60 sec bins) for animals in the threat (red) and control (blue) groups during the final training (pre-test), baseline (pre-stimulus) and stimulus exposure (stimulus) sessions. Threat exposure induced a significant (**D**) increase in time spent in the retreat box, (**E**) decrease in exploration, (**F**) increase in lurking, and (**G**) increase in staring, but no significant (**H**) change in scanning (mean of first 5 minutes; N = 5; ****P < 0.0001, **P < 0.01, *P < 0.05).

**Figure 2.**
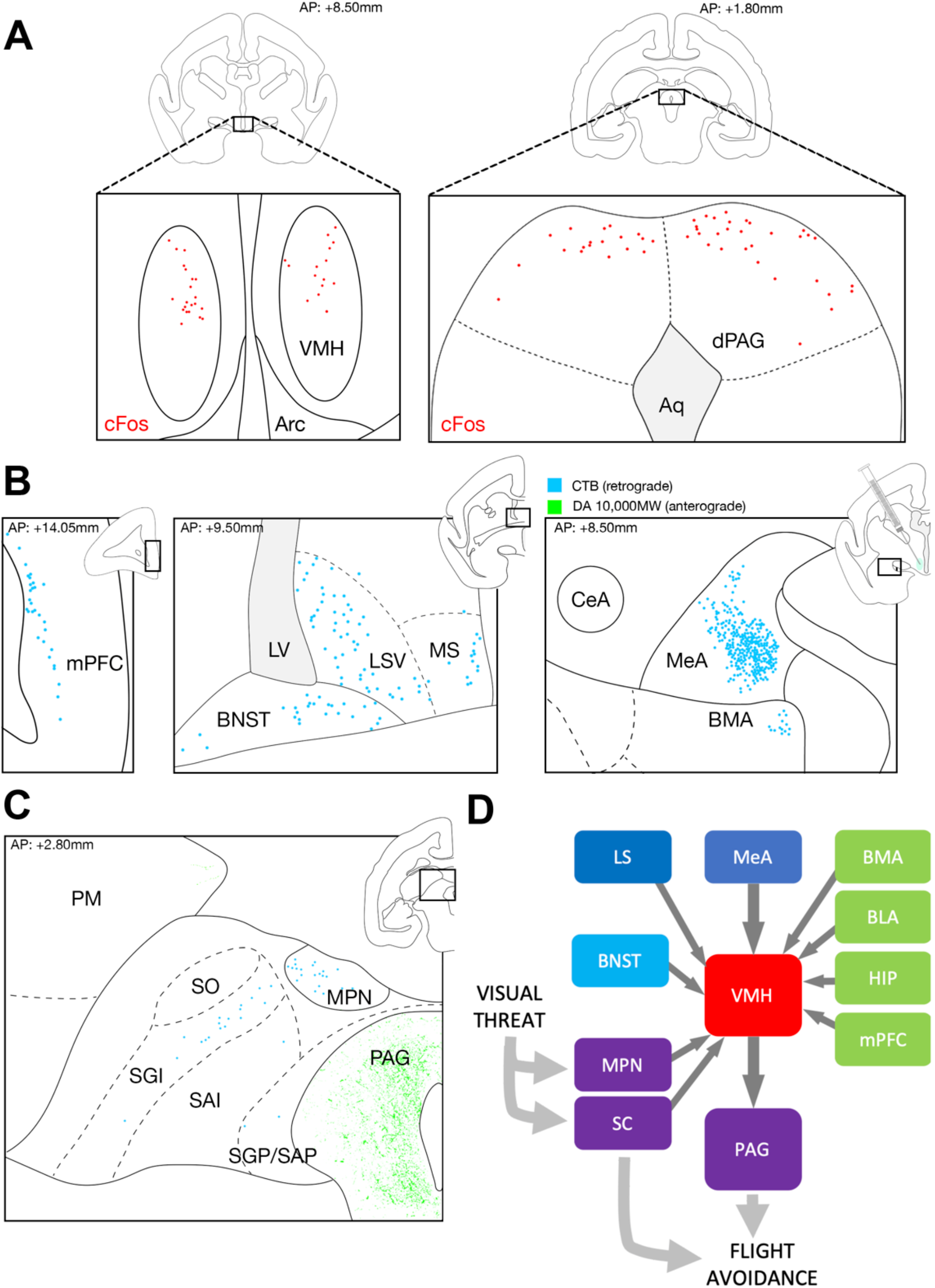
Threat exposure recruits the primate medial hypothalamic defensive system. (**A**) Robust cFos immunostained cells were found in the dorsomedial VMH and dorsalmedial PAG in animals exposed to snake threat, but not those exposed to the control stimulus (red dots indicate cFos+ cells in representative animal; N = 2). (**B**) Retrograde and anterograde tracing revealed sparse VMH afferents from mPFC, BNST, LS, MS, BMA and BLA and dense inputs from MeA. (**C**) Dense VMH efferents were found in PAG and sparse outputs in PM. (**D**) Summary of the VMH afferents and efferents in the primate. Dark grey arrows denote pathways that were identified in this study.

### Threat exposure recruits the primate medial hypothalamic defensive system

To determine whether the medial hypothalamus and its efferent and afferent targets are recruited in primates following predator exposure, we performed immunostaining for cFos – an immediate-early gene, in brain sections from experimental and control animals that had previously undergone injection of anterograde and retrograde tracers unilaterally into VMH (**Fig. 2B, right**). Within VMH, cFos+ cells were detected in the dorsomedial division (VMHdm), the subnucleus known to be essential for predator defense in rodents and whose activation is sufficient for the induction of fear and panic in humans (11) (**Fig. 2A left**, **Fig. S3A,C**). Importantly, no cFos+ cells were detected in VMH of control animals (**Fig. S3B**) or in the ventrolateral division of VMH (VMHvl) known to mediate defensive responses to social threat (8,19). Consistent with similar studies in rodents (20) cFos+ cells were also detected prominently in experimental, but not control animals in the dorsal periaqueductal grey (dPAG; **Fig. 2A right**, **Fig. S4**), a region known to be required for expression of predator defense in rodents (21,22). Finally, we identified cFos+ cells in the arcuate and paraventricular hypothalamic nuclei (**Fig. S3AB**), although similar numbers of immunopositive cells were seen in experimental and control animals, suggesting generalized recruitment of these structures during behavioral testing. Due to the high immunoreactive background, we were unable to determine the extent of cFos labeling in forebrain areas such as amygdala or medial prefrontal cortex (mPFC).

Anterograde and retrograde tracer delivery were restricted to VMH in three of four animals, allowing us to identify afferent and efferent regions of this structure in the primate brain and compare them with similar studies in the rodent. Sparse retrograde tracer-labeled cell bodies were found in the ventral mPFC (**Fig 2B left**, **Fig. S5AB**), medial division of the bed nucleus of the stria terminalis (BNST; **Fig 2B middle**, **Fig. S5C**), ventral division of the medial and lateral septum (respectively MS and LS; **Fig 2B middle**, **Fig. S5C**), posterior basomedial amygdala (BMA; **Fig. 2B right**, **Fig. S6A,CD**), and basolateral amygdala (BLA; **Fig. S6A,B,D**). Dense retrograde tracer-labeled cell bodies were found in the ventral division of the medial amygdala (MeA; **Fig. 2B right**; **Fig. S6A,D,E**), consistent with this structure providing significant inputs to VMH in rodents (23). Major anterograde tracer-labeled axonal varicosities, on the other hand, were found in the periaqueductal grey (PAG; **Fig. 2C**, **Fig. S7AB,F**) and sparse anterograde label was found in the medial pulvinar (PM; **Fig. 2C**, **Fig. S7A-C)**, intermediate layer of the superior colliculus (SGI; **Fig. 2C**, **Fig. S7A-B,D**) and medial pretectal nucleus (MPN; **Fig. 2C**, **Fig. S7 AB,E**). A few anterograde varicosities were also visible in MeA, BNST, and LS, but these could not be reliably confirmed.

## Discussion

The mechanism by which the amygdala is recruited and how it might influence defensive behavior remains contested (1,24). It is proposed that visually-evoked defensive responses to predators depends on fast, brainstem information processing via retino-collicular projections (24,25). From there, threat information passes to mesencephalic motor initiation centers to drive fixed medullary motor programs (26,27). The importance of this pathway is confirmed by the observation that SC lesions in primates abrogate both orienting and anxiety responses to a predator (28,29). At the same time, amygdala lesions also block fear and anxiety responses to predators in primates and humans (3,4). Here we show that the medial hypothalamic defensive system, known to be essential for defensive responses to predator in rodents (6–8), is engaged during predator defense in the marmoset.

This finding has two important implications for our understanding of fear in humans. First, our discovery makes it likely that the medial hypothalamus plays a similar role in encoding an internal state of threat in primates as it does in rodents (10,21,30–32). Notably, VMHdm in rodents is required for the induction and expression of both innate and conditioned predator fear (8,10,21) and stimulation of VMHdm in monkeys and humans is sufficient to elicit an intense defensive emotional state (11,33). The conserved recruitment of medial hypothalamus across species means that these structures must be considered in the search for the etiology and therapeutic treatment of anxiety-related disorders in humans.

Second, our identification of conserved connectivity of the medial hypothalamic defensive system across rodents and primates supports a common function for this system in defensive behaviors. In particular, the discovery that the primate VMH also receives major inputs from MeA, a nucleus known to convey information from the accessory olfactory system in rodents (34,35), was unexpected as this sensory system is vestigial in primates (36). These results suggest that MeA may have elaborated its non-olfactory inputs as vision evolved to become the dominant sense in primates. Finally, the existence of inputs to VMH from SC and MPN offers a direct pathway for visual information to rapidly enter the medial hypothalamic defensive system (**Fig. 2D**) (34,37) Nevertheless, given that amygdala lesions block fear responses to predators in humans 3, a parallel, indirect route that brings visual information to forebrain structures and from there to the medial hypothalamus is likely to also be important for both the emotional and behavioral responses to visual threat. In summary, our data argue for a conserved role of medial hypothalamic instinctive behavior networks across mammals, including humans. Further work is required to assess precisely which aspects of defensive behavior they control and whether they are necessary for generating the conscious emotional states that accompany threat in humans.

## Materials & Methods

### Animals

Six Common Marmosets (*Callithrix jacchus*) aged 18-24 months were sourced from the Australian National Nonhuman Primate Breeding and Research Facility. All experiments were conducted in accordance with the Australian Code of Practice for the Care and Use of Animals for Scientific Purposes and were approved by the Monash University Animal Ethics Committee, which also monitored the welfare of these animals.

### Behavioral assay

Testing was conducted in a transparent plexiglass transport box (305 × 295 × 205 mm) connected by a circular opening to a detachable black plexiglass nest box (600 × 140 × 130 mm) and placed on a table in the testing room in front of a removable black cloth canopy that covered the threat and control stimuli, respectively, a black striped rubber toy snake and a square black cardboard box. The snake could be animated by manipulating a pulley system from outside the isolated experimental room. On the day of the stimulation, a separate pulley system was used to lift the canopy, rapidly revealing the stimulus without the need for the experimenter to enter the room. A camera (GoPro Hero4) was positioned in front of the transport box to record the animal’s behavior. Manipulation of the stimulus and monitoring of the animal were all conducted in a room separate from the animal’s experimental room. The experimenter remained out of sight of the animals in a different room during testing. All animals underwent initial training for habituation to the experimental room. During this period, the animals were trained to enter the transport box that had been mounted to their home cage. Once the animals were habituated to the transport box, they were transported daily to the experimental room for habituation. There, the transport box was connected to the nest box, and the animals were allowed to explore the apparatus freely. The duration of habituation sessions was gradually increased from a few minutes to 20 minutes until the animal remained calm and relaxed during the entire session. At the end of each training session, a reward was given according to the animal’s preference before being brought back to their home cage. All training sessions commenced at 15:00 hours. Total time of transport box training and experimental room habituation was approximately two months. Once all animals were similarly habituated, four animals underwent tracer injection surgery and, following seven days of recovery, an additional 5 days of experimental room habituation to reinforce the training (20 minutes per session). Following retraining of the animals were exposed to either the control (N = 2) or threat stimulus (N = 2). Optimization of the control and threat stimulus presentation was conducted with the animal that did not undergo tracer surgery. Experimental validation involved testing behavioral responses to the threat and control stimuli and assessing recovery (N = 3, 5 minutes each: free apparatus exploration, stimulus exposure, stimulus occlusion, free apparatus exploration). The animals were randomly assigned to threat (N = 2, one male, one female) or control (N = 2, one male, one female) stimulus conditions (5 minutes free exploration, 45 minutes stimulus exposure). Following testing, the animal was rapidly processed for histological analysis.

### Behavior assessment

Videos were scored offline using BORIS software (38). The experimental space was divided into 6 zones: close vs far from the stimulus, standing vs close to the ground, and head out vs head in while in the retreat box. A weighted Defensive Behavior Index was calculated from these scores and represented as a continuously varying measure or heatmap. Exploration was estimated by counting the passage of the animal between zones. For several measures, data from both groups of animals were included as the protocols were indistinguishable over the first ten minutes of testing. Lurking was considered as both peeking (body in the retreat box, head out) and body and head in the retreat box while keeping the threat in sight. Staring was scored by counting each time the animal looked at the stimulation area. Scanning was scored by counting each time the animal looked upwards and around. All behaviors were analyzed by unpaired t-test.

### Surgery

Preparation of the animals for MRI-guided microinjection of the VMH was performed as previously described (18). In brief, animals were anaesthetized and scanned in a 9.4 T small-bore animal scanner. To facilitate reconstruction of the marmoset brain and visualization of the VMH structural T2 images were acquired with parameters included the following – repetition time/echo time: 6,000/40 ms, echo train length: 4, field of view: 38.4 × 38.4 mm^2^, acquisition matrix: 192 × 192, 100 coronal slices adjusted according to the size of the brain, slice thickness: 0.4 mm, signal averages: 4, scan time: 19 min, 42 s. Subsequently, the left hemisphere VMH was pressure injected with 180 nl of a bi-directional neural tracer cocktail consisting of 5 μg/μl retrograde Cholera toxin subunit B conjugated with Alexa Fluor 488 (Life Technologies, cat# C22841) and 50 μg/μl anterograde dextran amine 10,000 MW conjugated with Alexa Fluor 488 (Life Technologies, cat# D22910). Animals were allowed one week to recover to facilitate transport of neural tracers. Following seven days of recovery, animals underwent behavioral testing.

### Tissue processing

At the conclusion of behavioral testing, animals were deeply anaesthetized with 100 mg/kg sodium pentobarbitone. Following apnea, animals were transcardially perfused with heparinized saline and 4% paraformaldehyde in 0.01M PBS. Brains were post-fixed for 24 hrs in 4% PFA before being serially dehydrated in sucrose (10%, 20%, and 30%) solutions before being snap-frozen in −50 C isopentane and cryosectioned in the coronal plane at 50 μm. Sections were divided into four series and stored free-floating in a cryoprotective solution consisting of 50% phosphate buffered saline (PBS), 30% ethylene glycol and 20% glycerol at −20 C. For each subject, a full series was mounted onto glass slides (Superfrost plus), dehydrated in serial alcohols and cleared in xylene before being mounted in DPX for analysis of tracer label.

### Histology and immunohistochemistry

Tissue sections were rinsed in PBS before undergoing pre-treatment in a blocking solution consisting of PBS with 10% normal donkey serum and 2% Triton-X for 1 h at room temperature. Following blocking sections were incubated with primary antibody in pre-treatment solution for 16-18 hrs at 4 C. Sections were then washed in PBS before incubation in donkey anti-rabbit Alexa Fluor 594 secondary antibody (1:1,000, Life Technologies, cat# A11058) for 1 h at room temperature and washed and counterstained with Hoechst (Life Technologies, cat# H3569). Primary antibodies used in this study were rabbit anti-cFos (1:1,000, Sigma Aldrich, cat# F7799) to assess neural activity and rabbit anti-parvalbumin (1:2,000, Swant cat# PV27) to delineate boundaries of the superior colliculus. Acetylcholinesterase staining was performed to allow for demarcation of amygdala subnuclei and layers of the superior colliculus. The staining protocol was adapted from previous studies(39).

### Microscopy and image processing

Imaging was performed on an Axio Imager Z1 microscope (Zeiss). Images were acquired with a Zeiss Axiocam HRm digital camera using Axiovision software (v. 4.8.1.0). The objectives used were Zeiss EC-Plan Neofluar 5×0.16, #420330-9901, EC-Plan Neofluar 10×0.3, #420340-9901; Plan Apochromat 20×0.8 #420650-9901; EC Plan Neofluar 40×1.3 oil #420462-9900. Filter sets used were Zeiss DAPI #488049-9901-000, Zeiss HE eGFP #489038-9901-000 and Zeiss HQ TR #000000-1114-462. Stitching of images and adjustments to contrast and brightness were performed using Adobe Photoshop CC2015. The line art, boundaries and contours for all figures were executed using Adobe Illustrator CC2015. Demarcation of areas was achieved with AChE and parvalbumin labeling and compared with the Marmoset Brain in Stereotaxic Coordinates(40).

## General

We thank Claire Warner for launching the project and helping to delineate regions of the marmoset VMH and MeA.

## Funding

The work was supported by EMBL and Monash University, the European Research Council (ERC) Advanced Grant COREFEAR to C.T.G. and J.A.B. is supported by a Senior Research Fellowship support (APP1077677) from the National Health and Medical Research Council (NHMRC), and L.W. is supported by the International Partnership Program of Chinese Academy of Sciences (172644KYS820170004) and National Science Foundation of China (91732304).

## Author contributions

All the behavioral experiments and their analysis were carried out by Q.M. Surgical experiments were performed by W.K. and J.A.B. Histological experiments and analysis were carried out by W.K and Q.M. C.T.G. and J.A.B. conceived the project and with Q.M. and W.K. designed the experiments. C.T.G., J.A.B., Q.M and W.K. wrote the manuscript.

## Competing interests

There are no competing interests.

---

**Figure S1.**
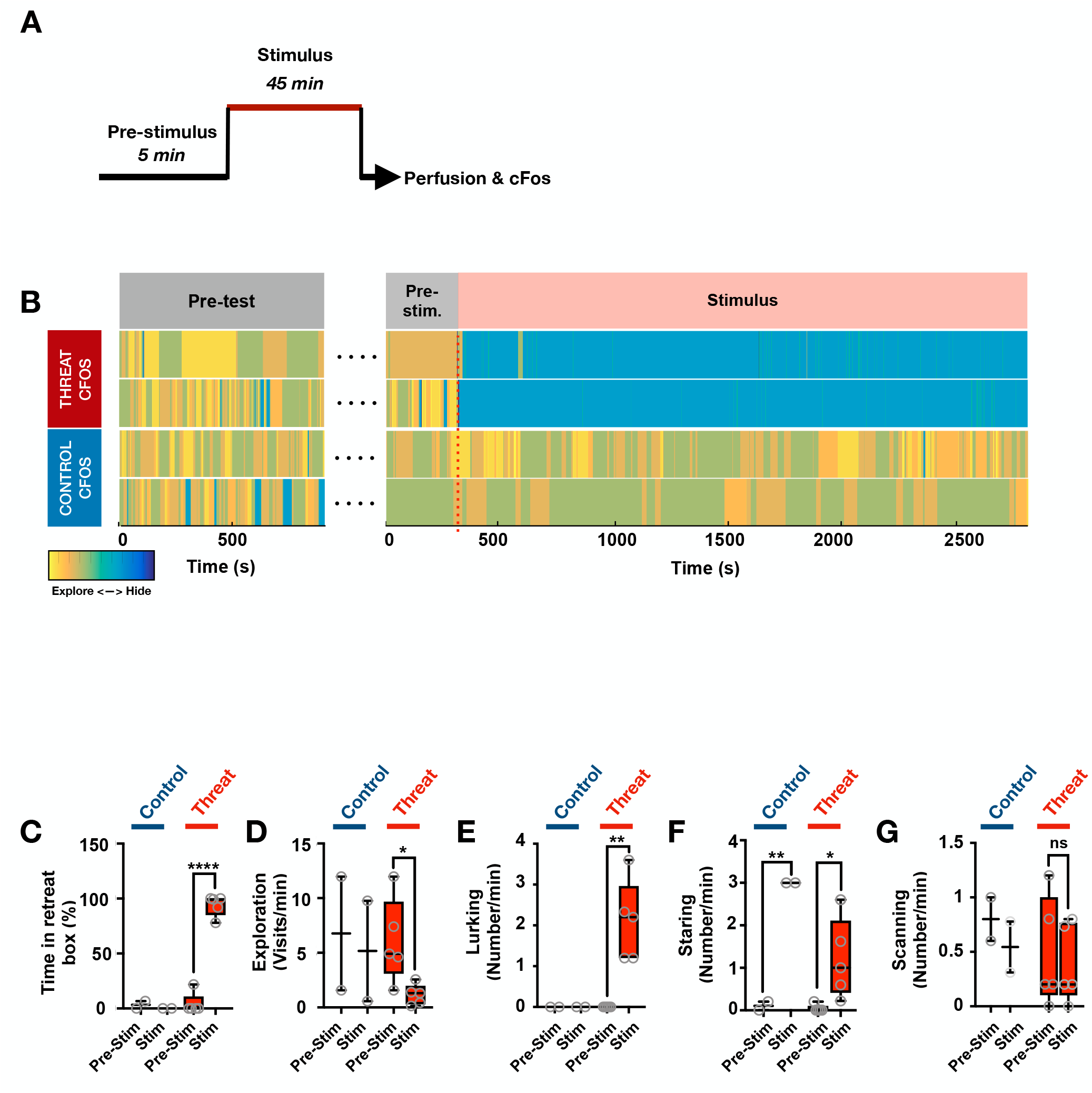
Behavior of animals in the cFos experiment. (**A**) Experimental phases: 5 min pre-stimulus during which the animal was freely exploring the apparatus, 45 min stimulus when the animal was exposed to the threat (toy animated snake) or control (black box) stimulus by raising a black cloth canopy, and post-experiment processing where the animal was anesthetized and perfused for histology and cFos immunistaining. (**B**) Defensive behavior for each animal in the threat (top) and control (bottom) groups. Color code indicates the animal is hiding in the retreat box (cold) or exploring (warm). (**C-G**) Quantification of behaviors of threat (red) and control (blue) groups during the pre-stimulus vs stimulus periods (5 min bins, N = 2-5; ****P < 0.0001, **P < 0.01, *P < 0.05).

**Figure S2.**
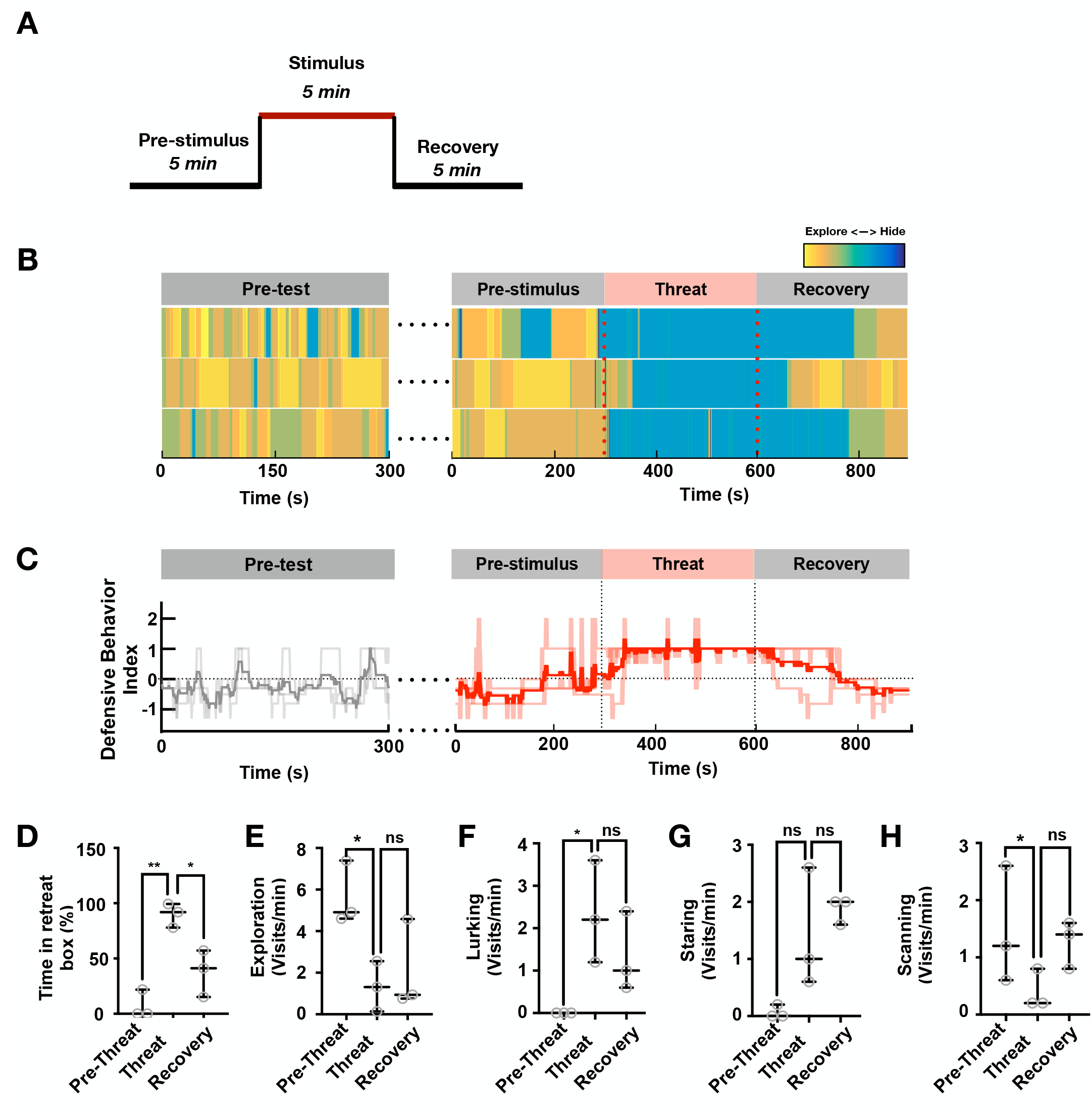
Behavior of animals in the recovery experiment. (**A**) Experimental phases: 5 min pre-stimulus during which the animal was freely exploring the apparatus, 5 min stimulus when the animal was exposed to the threat (toy animated snake) stimulus by raising a black cloth canopy, and recovery period when the stimulus was covered again by the cloth canopy. (**B**) Defensive behavior for each animal. Color code indicates the animal is hiding in the retreat box (cold) or exploring (warm). (**C**) Defensive Behavior Index (light color, individual traces; dark color, mean; 60 sec bins) for animal during the pre-test, pre-stimulus, threat, and recovery phases. (**D-H**) Quantification of behaviors during the pre-threat, threat, and recovery phases (N = 3; **P < 0.01, *P < 0.05).

**Figure S3.**
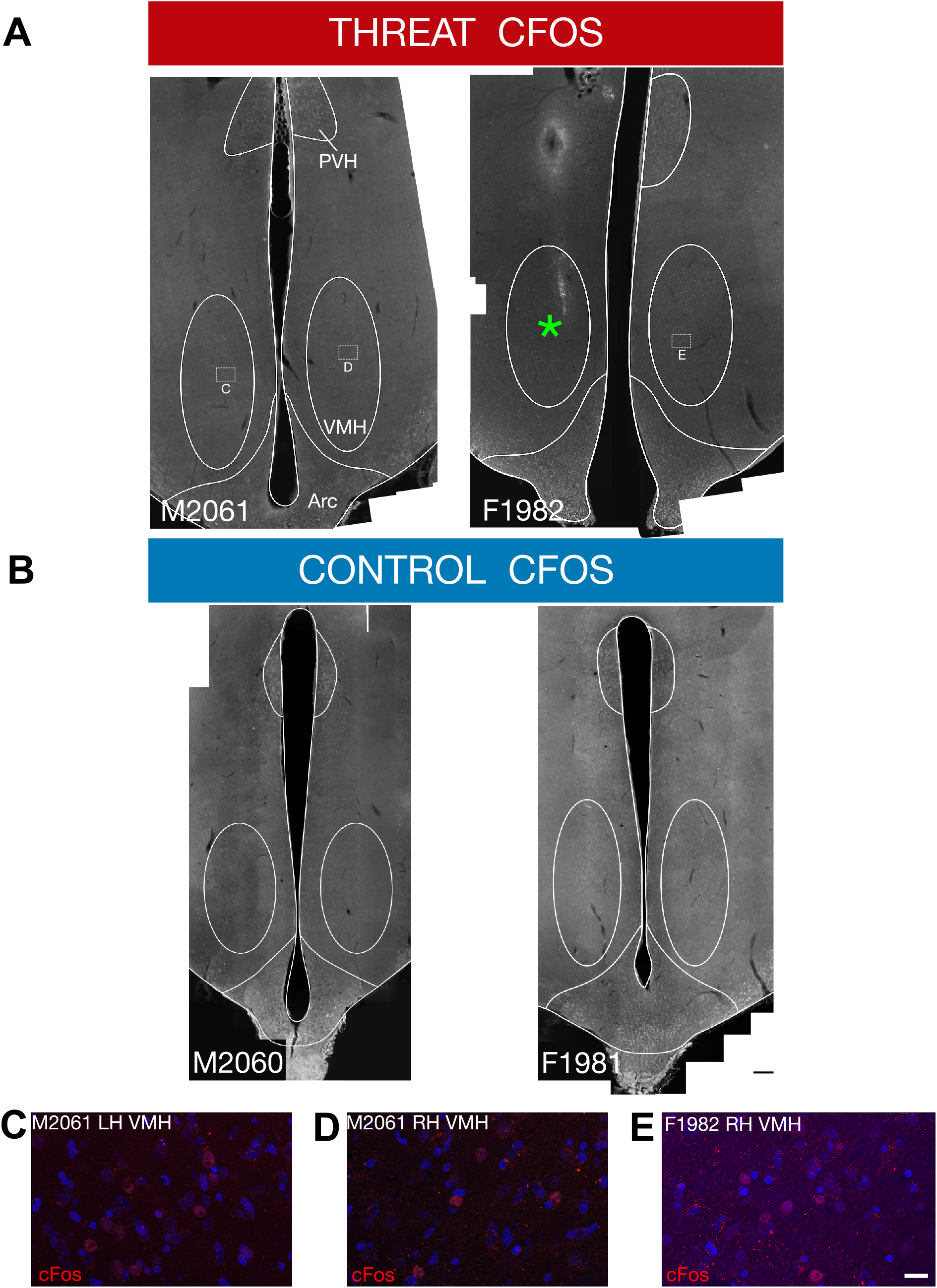
Threat-evoked cFos immunolabel in VMH. Representative coronal sections of the marmoset brain showing cFos immunolabeling in VMH of animals exposed to the (**A**) threat or (**B**) control stimulus. Green asterisk denotes needle tract from tracer administration. (**C-E**) High powered images from insets showing cFos+ cells identified within the VMH. No cFos+ cells were identified in animals exposed to the control stimulus. The arcuate (Arc) and paraventricular nucleus of the hypothalamus (PVH) contained cFos+ cells in both threat and control animals (numbers indicate animal ID; A, B: scale = 200 μm; C-E: scale = 20μm).

**Figure S4.**
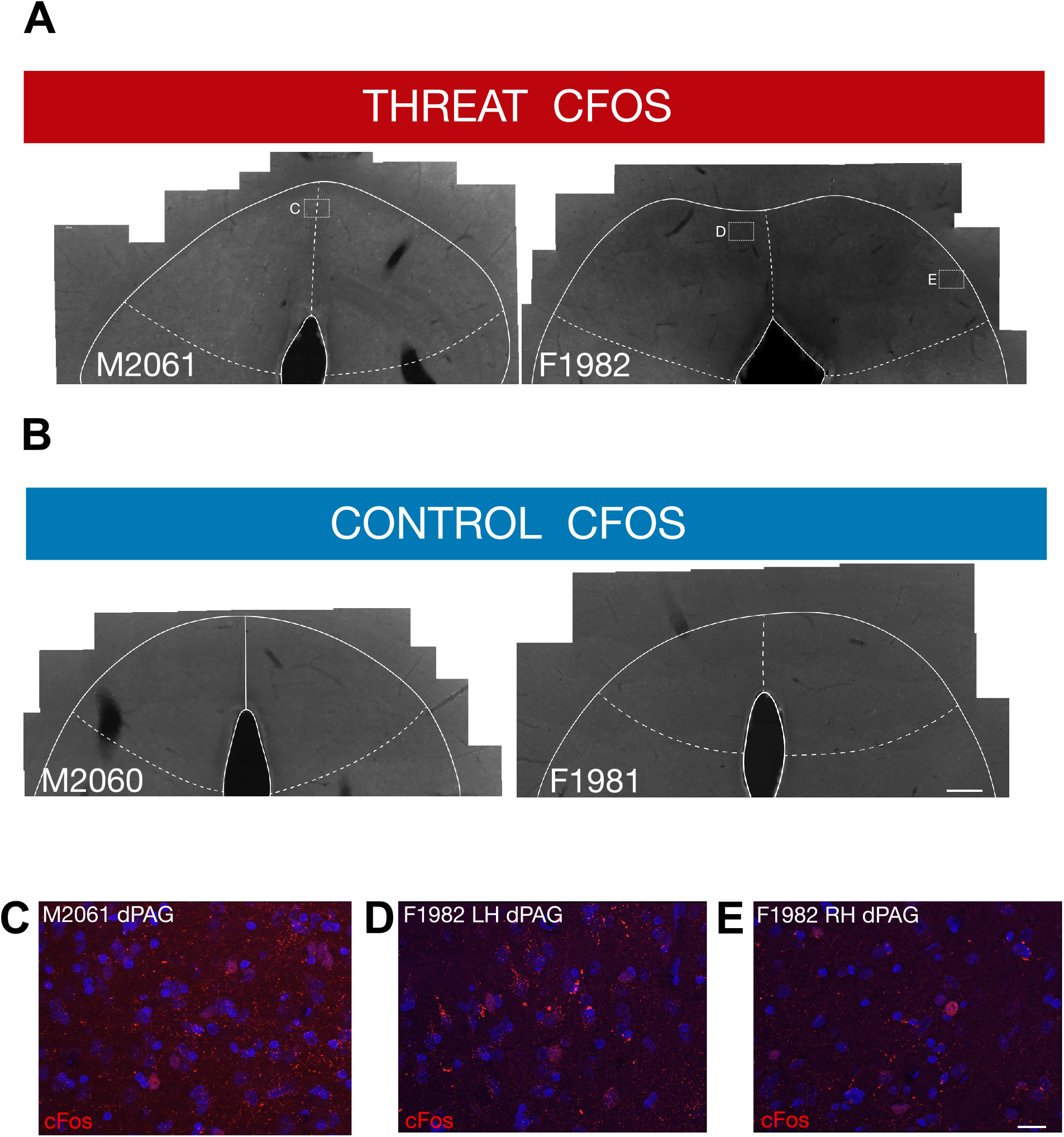
Threat-evoked cFos immunolabel in PAG. Representative coronal sections of the marmoset brain showing cFos immunolabeling in dPAG of animals exposed to the (**A**) threat or (**B**) control stimulus. (**C-E**) High powered images from insets showing cFos+ cells identified within the dPAG. No cFos+ cells were identified in animals exposed to the control stimulus (numbers indicate animal ID; A, B: scale = 200 μm; C-E: scale = 20μm).

**Figure S5.**
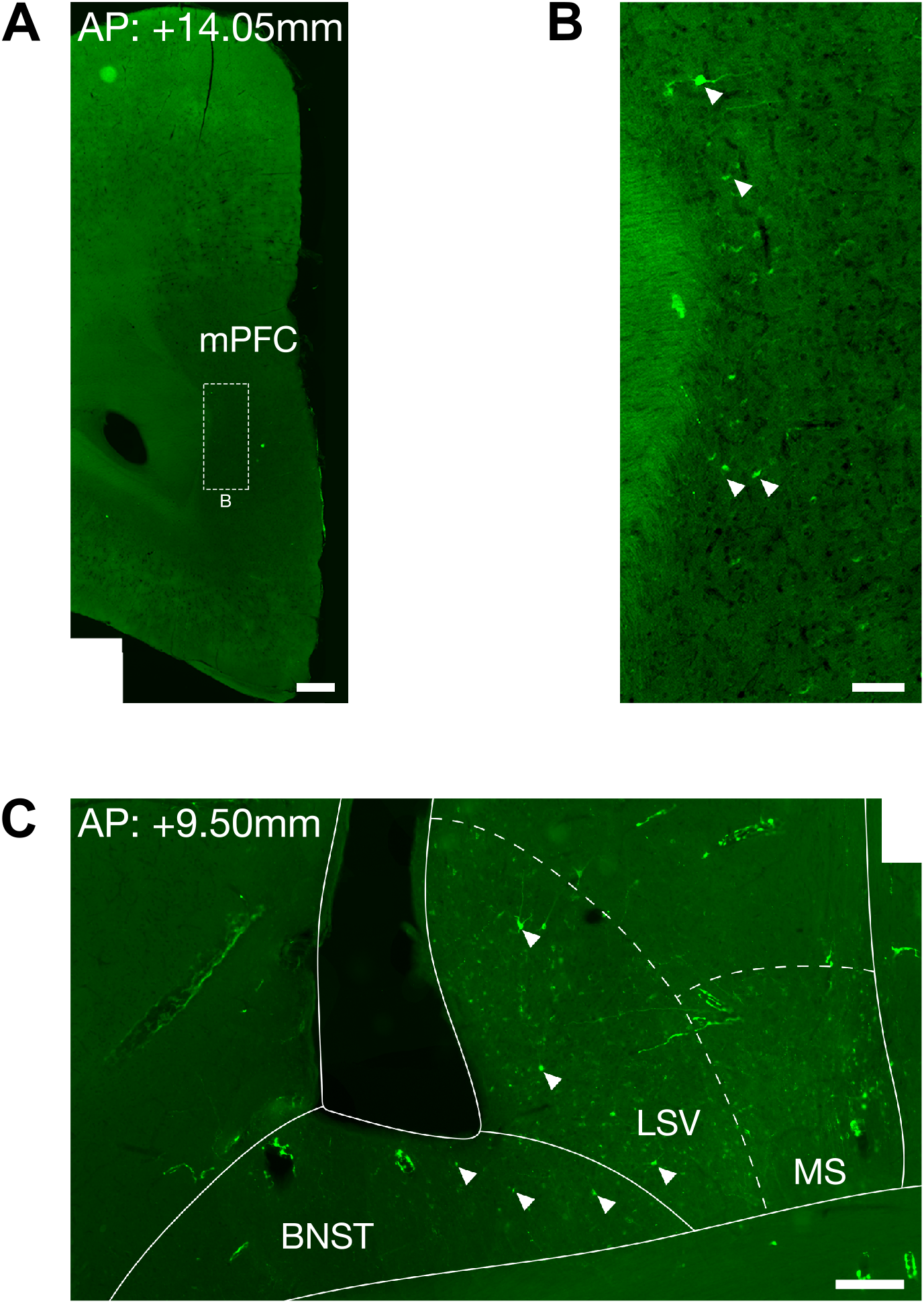
VMH connectivity in the forebrain. Representative coronal sections of the marmoset brain into which retrograde and anterograde tracers were delivered into VMH showing retrograde labeled cell bodies in (**A**, **B** high resolution inset) medial prefrontal cortex (mPFC) and (**C**) bed nucleus of the stria terminalis (BNST) and lateral and medial septum (LS and MS, respectively; scale: A 500 μm, B 100 μm, C 200 μm).

**Figure S6.**
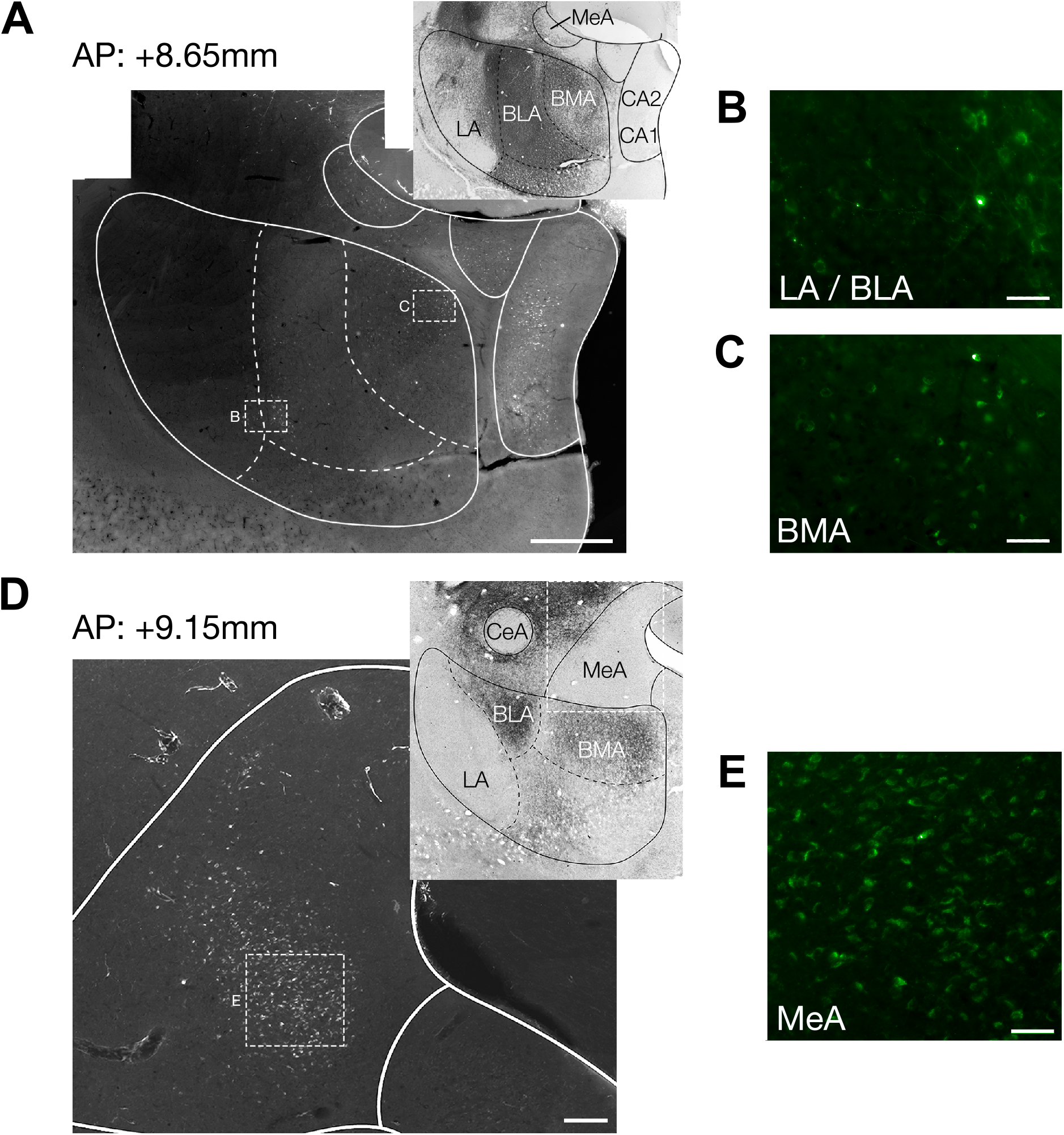
VMH connectivity in amygdala. Representative coronal sections of the marmoset brain into which retrograde and anterograde tracers were delivered into VMH showing retrograde labeled cell bodies in (**A**, high resolution insets **B-C**) BLA and BMA and (**D,** high resolution inset **E**) MeA (grayscale inset shows acetylcholinesterase stained adjacent section that was used to delineate brain regions; scale: A 1000 μm, B-C and E 50 μm, D 100μm).

**Figure S7.**
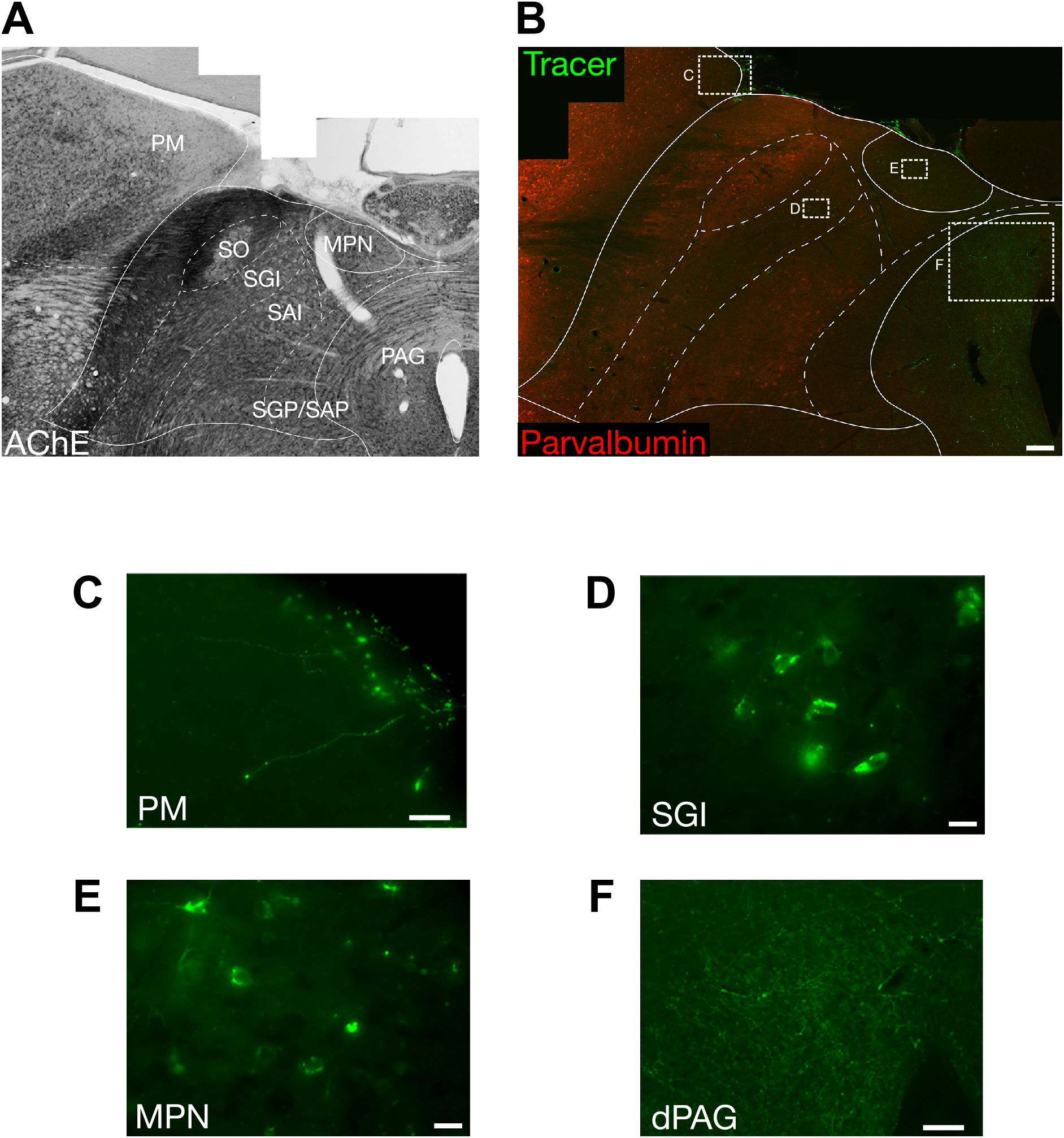
VMH connectivity in midbrain and thalamus. Representative coronal sections of the marmoset brain into which retrograde and anterograde tracers were delivered into VMH showing anterograde labeled processes in (**A-B** low resolution images with insets **C-F** ; AChE and Parvalbumin to delineate anatomical boundaries) PAG and PM, and retrograde labeled cell bodies in the SGI and MPN (scale: C 50 μm, D-E 20 μm and F 100μm).

## Supplementary Materials

Fig. S1. Behavior of animals in the cFos experiment.

Fig. S2. Behavior of animals in the recovery experiment.

Fig. S3. Threat-evoked cFos immunolabel in VMH.

Fig. S4. Threat-evoked cFos immunolabel in PAG.

Fig. S5. VMH connectivity in the forebrain.

Fig. S6. VMH connectivity in the amygdala.

Fig. S7. VMH connectivity in the midbrain and thalamus.

Movie S1. Animal response to threat stimulation.

Movie S2. Animal response to neutral stimulation.

## Notes

### Competing Interest Statement

The authors have declared no competing interest.

